# Serum response factor is essential for synaptic maturation in the hippocampus

**DOI:** 10.1101/2020.12.10.417360

**Authors:** Anna Krysiak, Matylda Roszkowska, Lena Majchrowicz, Anna Beroun, Piotr Michaluk, Karolina Nader, Martyna Pekala, Jacek Jaworski, Ludwika Kondrakiewicz, Alicja Puścian, Ewelina Knapska, Leszek Kaczmarek, Katarzyna Kalita

**Author notes:** To whom correspondence should be addressed, Researcher ID: R-4072-2016.

## Abstract

Disturbances of gene expression patterns that occur during brain development can severely affect signal transmission, connectivity, and plasticity—key features that underlie memory formation and storage in neurons. Abnormalities at the molecular level can manifest as changes in the structural and functional plasticity of dendritic spines that harbor excitatory synapses. This can lead to such developmental neuropsychiatric conditions as Autism spectrum disorders, intellectual disabilities, and schizophrenia. The present study investigated the role of the major transcriptional regulator serum response factor (SRF) in synapse maturation and its impact on behavioral phenotypes. Using *in vitro* and *in vivo* models of early postnatal SRF deletion, we studied its influence on key morphological and physiological hallmarks of spine development. The elimination of SRF in developing neurons resulted in a phenotype of immature dendritic spines and impairments in excitatory transmission. Moreover, using a combination of molecular and imaging techniques, we showed that SRF-depleted neurons exhibited a lower level of specific glutamate receptor mRNAs and a decrease in their surface expression. Additionally, the early postnatal elimination of SRF in hippocampal CA1 excitatory neurons caused spine immaturity and a specific social deficit that is frequently observed in autism patients. Altogether, our data suggest that the regulation of structural and functional dendritic spine maturation begins at the stage of gene transcription, which underpins the crucial role of such transcription factors as SRF. Moreover, disturbances of the postnatal expression of SRF translate to behavioral changes in adult animals.

## Introduction

Neuronal development is controlled at the transcriptional level. A growing body of evidence shows that the development of neuropsychiatric diseases is linked to mutations of genes that encode proteins that control activity-dependent transcription and chromatin remodeling (Chen *et al*, 2017; Ebert & Greenberg, 2013; Hong *et al*, 2005; West & Greenberg, 2011; Yap & Greenberg, 2018). Dysregulation of the global gene expression program in the developing brain may result in alterations of the number of neuronal connections and synaptic strength and shape, leading to unbalanced neuronal plasticity (Aguado *et al*, 2009; Del Blanco *et al*, 2019; Flavell *et al*, 2006; Shalizi *et al*, 2006).

Most excitatory synapses are located on specialized subcellular compartments, called dendritic spines. Dendritic spines are morphologically heterogeneous protrusions that show a correlation between their structure and function (Holtmaat & Svoboda, 2009; Kasai *et al*, 2010; Kasai *et al*, 2003). During brain development and in response to an increase in neuronal activity, dendritic spine shapes change from immature, thin, elongated filopodia-like structures to stable, mature mushroom spines (Fiala *et al*, 1998; Hering & Sheng, 2001; Papa & Segal, 1996). Spine morphology is also tightly linked to the abundance of α-amino-3-hydroxy-5-methyl-4-isoxazolepropionic acid (AMPA) glutamate receptors and efficacy of excitatory synapses (Henley & Wilkinson, 2016; Nimchinsky *et al*, 2002). Structural spine plasticity is crucial for the plasticity of neuronal circuits and supports the formation and long-term storage of information in the brain (Holtmaat & Svoboda, 2009; Kasai *et al.*, 2010; Yang *et al*, 2009). Abnormalities in the number and shape of dendritic spines are frequently observed in developmental neuropsychiatric disorders, such as fragile X syndrome, autism spectrum disorder (ASD), schizophrenia, and intellectual disability (Glausier & Lewis, 2013; Levenga & Willemsen, 2012; Nimchinsky *et al*, 2001; Penzes *et al*, 2011; Phillips & Pozzo-Miller, 2015).

Serum response factor (SRF) is a major transcription factor in the brain. It regulates the activity-dependent expression of genes and encoding of proteins that are involved in synaptic plasticity (Kuzniewska *et al*, 2016; Kuzniewska *et al*, 2013; Losing *et al*, 2017; Parkitna *et al*, 2010; Ramanan *et al*, 2005). SRF is essential for proper neuronal development. Its global deletion causes death in mice during gastrulation because of problems in generation of the mesoderm (Arsenian *et al*, 1998). The availability of conditional mutant mice has allowed demonstrations that SRF controls neuronal cell migration, neurite outgrowth, organization of the dentate gyrus (DG), and the formation of mossy fiber circuitry during brain development (Alberti *et al*, 2005; Etkin *et al*, 2006; Knoll *et al*, 2006; Li *et al*, 2014; Lu & Ramanan, 2011; Scandaglia *et al*, 2015; Stritt & Knoll, 2010). Moreover, the early postnatal inactivation of SRF leads to deficiencies in long-term potentiation, long-term depression, learning, and memory (Etkin *et al.*, 2006; Ramanan *et al.*, 2005). Altogether, these observations suggest that SRF controls gene expression programs that are critical for neuronal connectivity and information processing in the brain. In contrast to the paramount role of SRF in brain development, its removal in adult excitatory neurons does not affect gross hippocampal architecture (Kuzniewska *et al.*, 2016; Losing *et al.*, 2017; Nader *et al*, 2019). Despite advances in understanding the role of SRF in the central nervous system, the contributions of SRF to dendritic spine formation and neurodevelopmental disorders remain unresolved.

The present study directly investigated the role of SRF in glutamatergic synapse maturation by deleting SRF in hippocampal pyramidal neurons *in vitro* and *in vivo*. We used mice that lacked SRF postnatally by crossing SRF^f/f^ mice with transgenic lines that expressed Cre recombinase (tamoxifen-regulated) under control of the CaMKIIα promoter to precisely investigate the role of SRF within the time window of spine maturation, starting from postnatal day 5-6 (P5-6). Our data showed that the loss of SRF in hippocampal neurons disrupted spine maturation at both the structural and functional levels. We also found that SRF deficiency downregulated expression of the AMPA receptor (AMPAR) subunits GluR1 and GluR2, leading to the formation of immature synapses. The lack of SRF during early stages of spine maturation led to an increase in the percentage of filopodial dendritic spines *in vivo* and alterations of specific aspects of social behaviors in SRF knockout (KO) mice.

## Results

### Depletion of SRF reduces the number of mature spines

To determine whether SRF-driven transcription in neurons might be developmentally regulated, we first analyzed SRF transcriptional activity *in vitro*. On DIV0, rat hippocampal neurons were electroporated using Amaxa with a 5×SRE reporter construct that contained five CArG boxes. SRF-driven transcription was analyzed on DIV3, DIV7, DIV14, and DIV21. An increase in transcription levels was observed on DIV7 to DIV21, with the highest level detected on DIV14 (**Fig. 1A**; DIV3: 1.00 ± 0.12, DIV7: 2.66 ± 0.29, DIV14: 3.76 ± 0.62, DIV21: 2.95 ± 0.26, *n* > 8, one-way ANOVA measures analysis of variance: *F*_3,31_ = 11.67; *p* < 0.0001, Sidak’s multiple-comparison *post hoc* test; *p* = 0.0040, DIV3 *vs*. DIV7; *p* < 0.0001, DIV3 *vs*. DIV14; *p* = 0.0011, DIV3 *vs*. DIV21). These results suggest that SRF-dependent transcription was developmentally regulated. Next, to assess the impact of SRF on synaptogenesis and dendritic spine maturation, we used the Lipofectamine-based transfection method, which allowed us to visualize only sparsely stained neurons and precisely analyze spine morphology. We first tested the efficiency of the shRNA-induced knockdown of SRF *in vitro* (**Fig. S1**). To decrease SRF expression, we expressed shRNA against SRF with GFP in hippocampal rat primary cultures on DIV6-7. As a control, scrambled shRNA (shCTR) was used. The cells were fixed on DIV18-21, followed by immunostaining against endogenous SRF. Comparisons of the average SRF immunofluorescence intensity showed a significant decrease in endogenous SRF protein expression at the single-cell level (**Fig. S1A**). Similarly, SRF deletion was evaluated in SRF^f/f^ mouse hippocampal neurons that were transfected with a plasmid that encoded Cre recombinase under the CamKIIα promoter (**Fig. S1B**). Next, we assessed the effects of SRF deficiency on the density and morphology of dendritic spines. The density analysis showed no significant changes in the number of dendritic protrusions on DIV21 between shCTR- and shSRF-transfected neurons, suggesting that SRF is not required for spine formation (**Fig. 1D**; shCTR: 0.53 ± 0.04, *n* = 22 cells; shSRF: 0.47 ± 0.04, *n* = 18 cells; Student’s *t*-test: *p* = 0.39). However, on DIV21, we observed a significant increase in dendritic spine length in SRF-depleted rat neurons (**Fig. 1E**; shCTR: 0.98 ± 0.02, *n* = 22 cells; shSRF: 1.30 ± 0.06, *n* = 18 cells; Student’s *t*-test: *p* < 0.001). To further investigate how the lack of SRF affects dendritic spine shape, we clustered spines into two distinct morphological categories: “filopodia and long spines” and “mushroom and stubby spines.” At a time point when the majority of developing excitatory postsynaptic structures exhibit a mushroom shape, we observed a significant shift in the spine morphology of SRF-depleted neurons to filopodia-like, immature protrusions compared with control cells. Therefore, although SRF deficiency did not affect the number of spines, the early developmental deletion of SRF significantly increased by ~24% the proportion of filopodia and long spines, with a significant decrease in the population of mushroom and stubby spines (**Fig. 1F**; shCTR, *n* = 22 cells; shSRF, *n* = 18 cells; *χ^2^* test, *df* = 11.99, 1, *p* = 0.0005) compared with shCTR neurons.

**Fig. 1.**
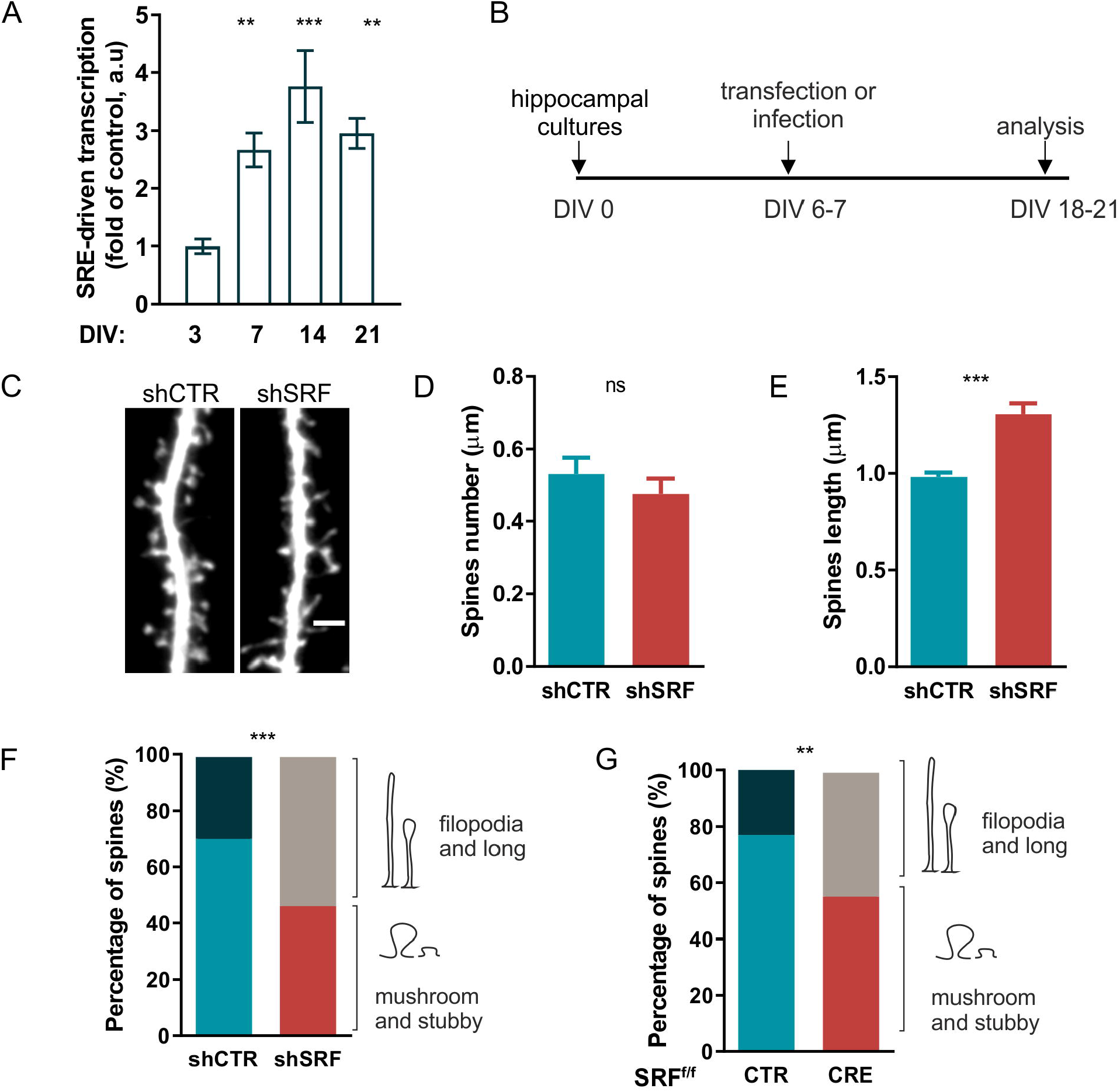
SRF regulates dendritic spine maturation. **(A)** SRF-driven transcription in rat primary hippocampal cultures on DIV3, DIV7, DIV14, and DIV21. Rat hippocampal neurons were co-transfected with a 5×SRE reporter together with β-Gal control plasmid on P0 (Amaxa). SRF-driven transcription was evaluated by determining the luciferase/β-Gal activity ratio. \*\**p* < 0.01, \*\*\**p* < 0.001 (one-way ANOVA followed by Sidak’s multiple-comparison *post hoc* test). The data are expressed as mean ± SEM. **(B)** Schematic diagram of the experimental design. **(C)** Example photographs of GFP-stained dendrites with dendritic spines of rat hippocampal neurons transfected with shCTR or shSRF plasmid. Scale bar = 2 μm. **(D, E)** Analysis of dendritic spine density and length in neurons that were transfected with shCTR and shSRF. **(F)** Percentage of protrusions that clustered into two categories (“filopodia and long spines” and “mushroom and stubby spines”) in rat hippocampal neurons that were transfected with shCTR or shSRF. **(G)** Percentage of protrusions that clustered into two categories in mouse hippocampal neurons that were transfected with a control vector (CTR) or CRE. *^ns^p* > 0.05, *\*\*p* < 0.01, *\*\*\*p* < 0.001 (Student’s *t*-test). The data were obtained from at least three independent neuronal cultures and are expressed as mean ± SEM.

To exclude the possibility that shSRF may act nonspecifically, we performed an independent analysis using mouse neurons that were isolated from the hippocampus of SRF^f/f^ mice that were transfected with a plasmid that encoded Cre recombinase under the CamKIIα promoter on DIV6-7. Similar to the shRNA results, mouse neurons that expressed Cre recombinase also exhibited a lower percentage of mature dendritic spines and an increase in immature protrusions (**Fig. 1G**; ~21% of filopodia and long spines AAV-CTR, *n* = 12; AAV-CRE, *n* = 17; *χ^2^* test, *df* = 10.24, 1, *p* = 0.0014), independently confirming that SRF facilitates structural spine maturation.

Morphological changes in the postsynaptic component of excitatory synapses have a potential to influence their function (Noguchi *et al*, 2011; Sala & Segal, 2014). Assuming that SRF is crucial for the structural maturation of dendritic spines, we next tested whether SRF deletion leads to functional alterations of excitatory synaptic transmission. Hippocampal neurons were transfected on DIV6 with either shSRF or shCTR. On DIV21, AMPAR-mediated miniature excitatory postsynaptic currents (AMPAR-mEPSCs) were recorded from GFP-expressing neurons.

Comparisons of the cumulative probability of cellular responses between shCTR- and shSRF-transfected neurons demonstrated a significant decrease in mEPSC amplitude (**Fig. 2B**; Kolmogorov-Smirnov test: *p* = 0.0080; **Fig. 2 C**, shCTR: 26.01 ± 0.96, *n* = 8 cells; shSRF: 22.9 ± 0.81, *n* = 10 cells, Mann-Whitney test, *p* = 0.0026) and frequency (**Fig. 2 D**; Kolmogorov-Smirnov test, *p* = 0.0002; **Fig. 2 E**, shCTR: 0.33 ± 0.040, *n* = 8 cells shSRF: 0.54 ± 0.047, *n* = 10 cells, Mann-Whitney test, *p* < 0.0001), indicating impairments in excitatory synaptic transmission. The increase in the time of inter-events intervals of mEPSCs compared with shCTR-transfected cells might indicate a lower number of mature synapses in SRF-depleted neurons.

**Fig. 2.**
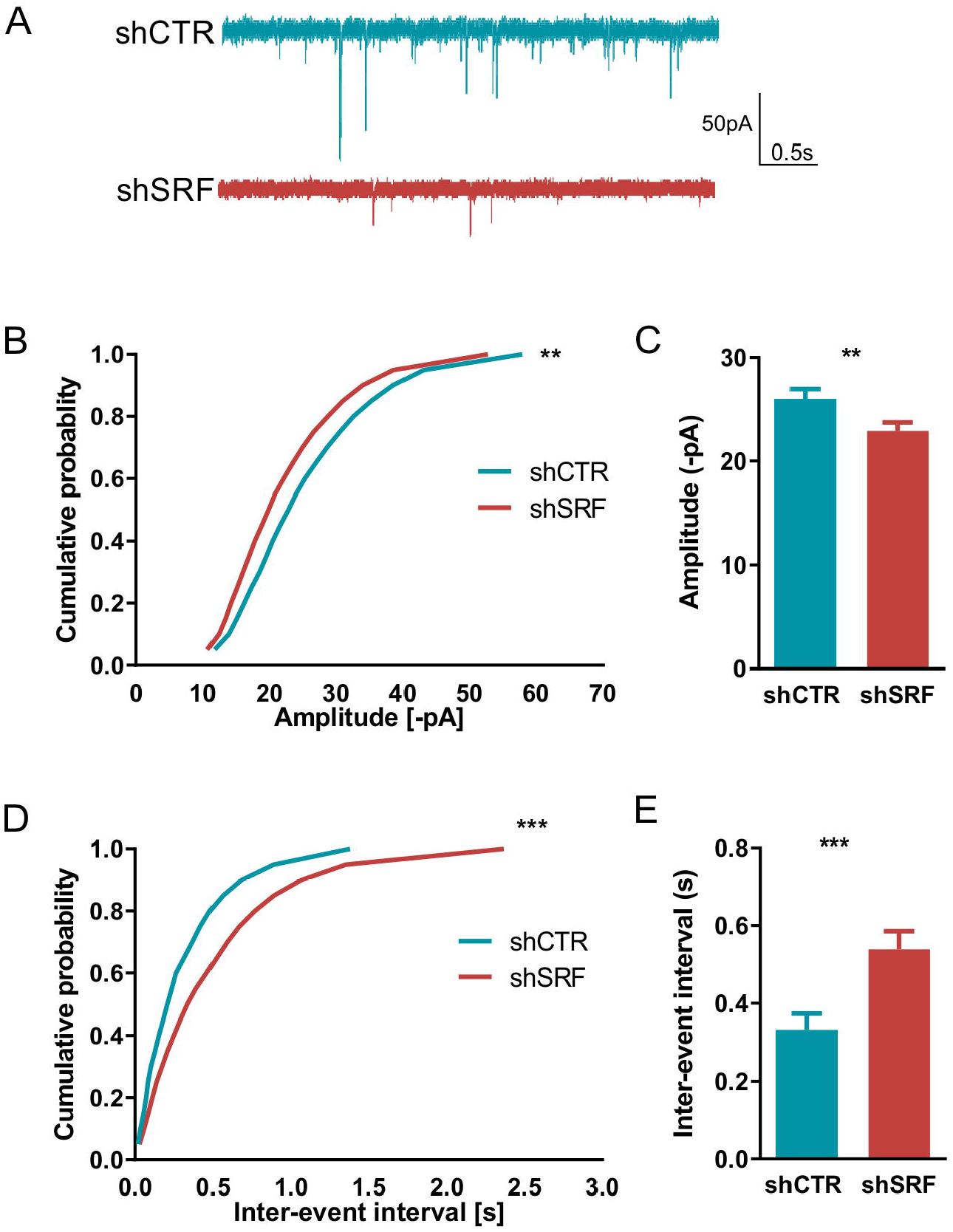
Lack of SRF decreases the amplitude and frequency of spontaneous mEPSCs in hippocampal neuronal cultures. **(A)** Sample traces of mEPSCs in shCTR- or shSRF-transfected hippocampal neurons. Cumulative probability plots show the mEPSC amplitude **(B)** and frequency **(D)** in shCTR cells and shSRF-transfected neurons from at least three independent neuronal cultures. *\*\*p* < 0.01, *\*\*\*p* < 0.001 (Kolmogorov-Smirnov test). Bar graphs in **(C)** and **(E)** represent average amplitudes and frequencies of mEPSCs. *\*\*p* < 0.01, *\*\*\*p* < 0.001 (Mann-Whitney test). The data are expressed as mean ± SEM.

### SRF regulates the number of functional synapses

We found that changes in spine morphology were associated with decreases in the amplitude and frequency of mEPSCs in SRF-depleted neurons. No differences in spine density were detected between the analyzed groups. Thus, the observed changes in synaptic transmission may have resulted from a postsynaptic defect. To test whether a lower a number of functional synapses underlies reduced mEPSC frequency, we quantified markers of the excitatory synapses as a morphological readout of neuronal connectivity. On DIV21, we measured the number of puncta that were stained with presynaptic (Bassoon) and postsynaptic (PSD-95) proteins in neurons that were transfected with either shCTR or shSRF (**Fig. 3**). The quantification of Bassoon-positive puncta density at selected dendritic segments revealed no significant changes between groups (**Fig. 3B**; shCTR: 1.02 ± 0.04, *n* = 34 cells; shSRF: 0.93 ± 0.04, *n* = 40 cells; Mann-Whitney test, *p* = 0.4105). Simultaneously, further analysis of the same dendrites revealed that shSRF-transfected neurons exhibited a decrease in PSD-95 puncta density (**Fig. 3C**; shCTR: 260.5 ± 13.02, *n* = 34 cells; shSRF: 151.7 ± 13.3, *n* = 40 cells; Mann-Whitney test, *p* < 0.0001) compared with control cells. Moreover, the immunofluorescence staining of primary hippocampal neurons revealed a decrease in the synaptic overlap of Bassoon and PSD-95 proteins in shSRF-transfected neurons, reflected by a lower Pearson’s correlation coefficient (**Fig. 3D**; shCTR, 0.54 ± 0.02, *n* = 32 cells; shSRF, 0.44 ± 0.02, *n* = 37 cells; Mann-Whitney test, *p* < 0.001). Altogether, these results suggest that the number of functional synapses decreased after silencing SRF expression during neuronal development *in vitro*.

**Fig. 3.**
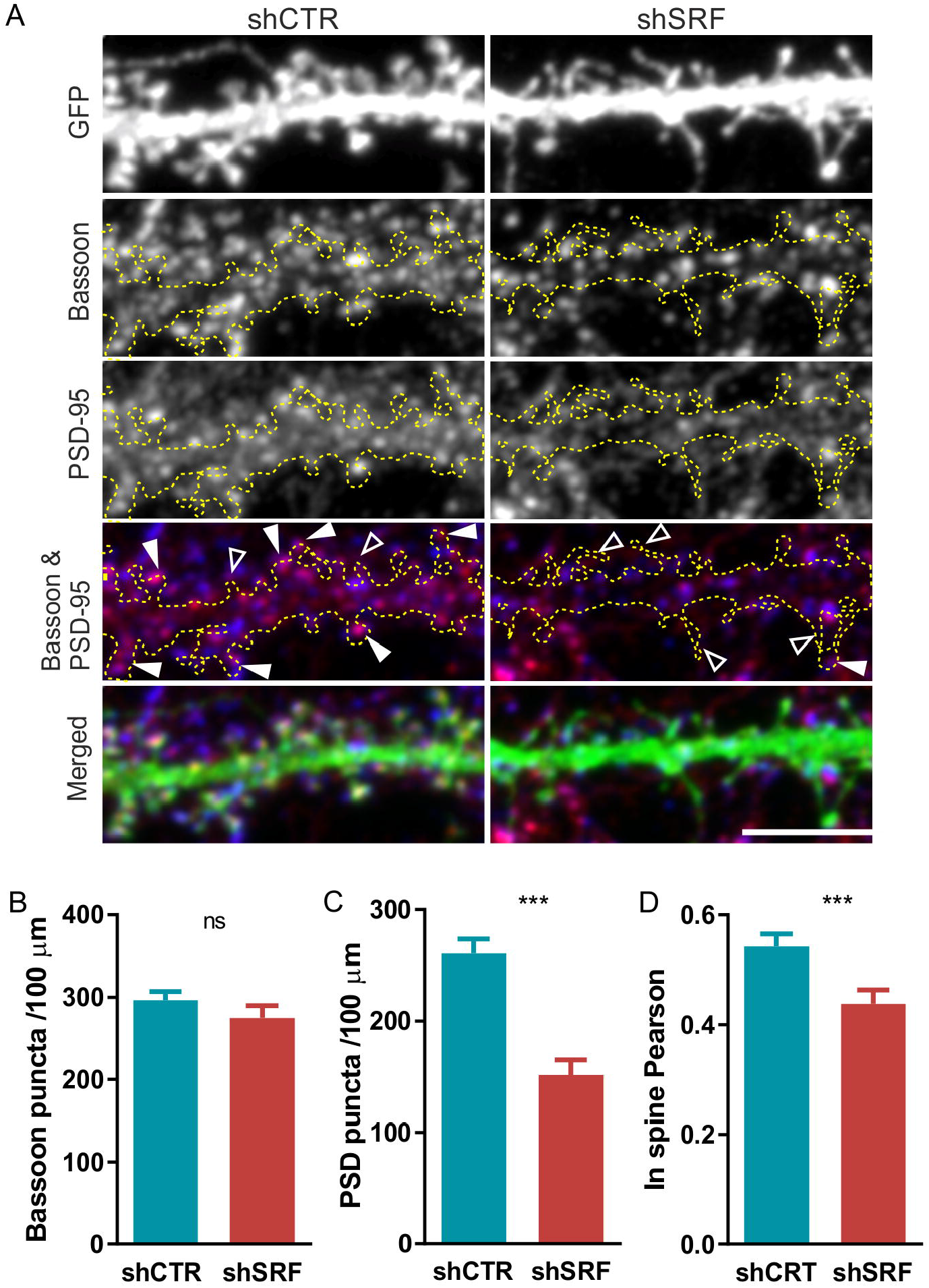
SRF regulates the number of functional synapses in hippocampal neurons. **(A)** Representative images of dendritic segments of shCTR- and shSRF-transfected neurons that were double-immunostained with anti-Bassoon (presynaptic protein; blue) and anti-PSD-95 (postsynaptic protein; red) antibodies. Dendrites of GFP-positive neurons were outlined. In the red/blue color scheme, the colocalization of both synaptic markers is shown as purple immunofluorescence. White arrowheads indicate an overlap of both synaptic markers within the dendritic spine. Empty arrowheads indicate Bassoon-positive/PSD-95-negative dendritic spines. Scale bar = 5 μm. **(B, C)** Quantification of Bassoon and PSD-95 puncta density within dendritic segments. **(D)** Overlap of Bassoon and PSD-95 puncta in dendritic spines, expressed as Pearson’s coefficient. The data were obtained from at least three separate experiments. *^ns^p* > 0.05, *\*\*\*p* < 0.001 (Mann-Whitney test). The data are expressed as mean ± SEM. Only Bassoon puncta density data **(B)** passed the normality test. ^ns^*p* = 0.4105 (Student’s *t*-test).

### SRF knockdown decreases surface levels of AMPARs

To further investigate the functional significance of the shift in spine morphology toward a more immature phenotype, we analyzed the impact of SRF deletion on surface AMPAR expression. To measure the level of surface-exposed GluR1 and GluR2 subunits, we used a live-staining method with antibodies that were directed against the extracellular N-terminal domains of GluR1 or GluR2 subtypes. The labeling procedure was performed in hippocampal neurons that were transfected with shCTR or shSRF (**Fig. 4**). SRF depletion significantly reduced the number of surface GluR1 (**Fig. 4B**; shCTR: 137.2 ± 22.43, *n* = 13 cells; shSRF: 70.48 ± 16.58, *n* = 14 cells; Mann-Whitney test: *p* = 0.0145) and GluR2 (**Fig. 4D**; shCTR: 142.2 ± 11.65, *n* = 13 cells; shSRF: 86.42 ± 17.96, *n* = 14 cells; Mann-Whitney test, *p* = 0.0017) puncta, suggesting a decrease in postsynaptic AMPAR levels, which was consistent with the reduction of mEPSC amplitude in shSRF-transfected neurons (**Fig. 2**). Altogether, these data demonstrate that SRF regulates AMPAR-dependent excitatory synaptic transmission.

**Fig 4.**
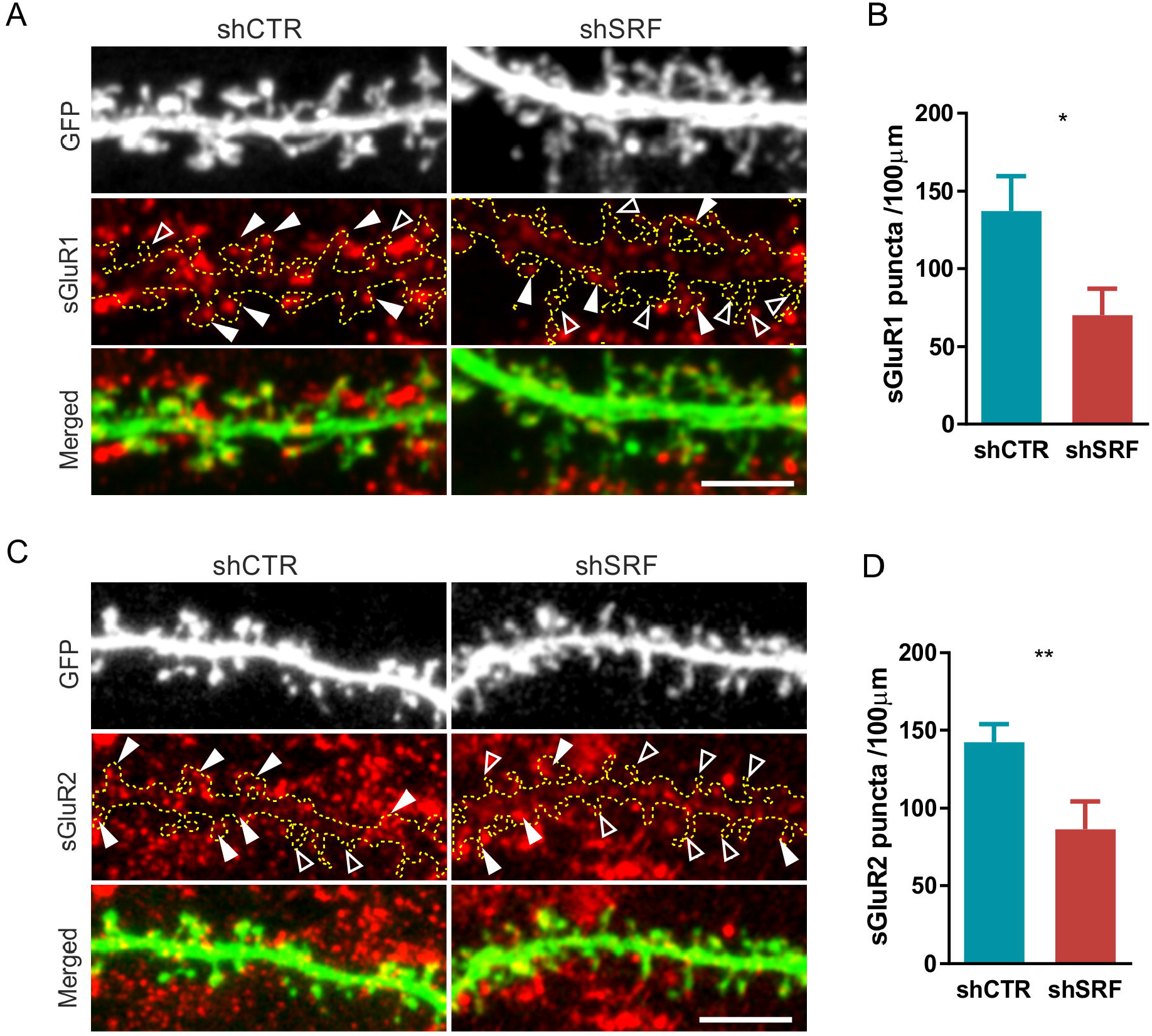
SRF depletion reduces surface GluR1 and GluR2 AMPAR subunit levels in rat hippocampal neurons. **(A)** Representative images of dendritic segments of shCTR- or shSRF-transfected neurons that were immunostained with anti-GluR1 antibody. Dendrites of GFP-positive neurons are outlined. Examples of GluR1-positive dendritic spines are indicated by white arrowheads. Examples of spines that lack GluR1 expression are indicated by empty arrowheads. Scale bar = 5 μm. **(B)** Quantification of GluR1 puncta density localized within the dendritic segment. **(C)** Representative images of dendritic segments of shCTR- or shSRF-transfected neurons that were immunostained with anti-GluR2 antibody. Dendrites of GFP-positive neurons are outlined. Examples of GluR2-positive dendritic spines are indicated by white arrowheads. Examples of spines that lack GluR2 expression are indicated by empty arrowheads. Scale bar = 5 μm. **(D)** Quantification of GluR2 puncta density localized at dendritic segments. The data were obtained from at least three separate experiments. \**p* < 0.05, *\*\*p* < 0.01 (Mann-Whitney test). The data are expressed as mean ± SEM.

To validate the results that showed the decrease in surface GluR1 and GluR2 levels in shSRF-transfected neurons, we performed a biochemical analysis of SRF-depleted rat hippocampal cultures. We first confirmed the efficiency of the AAV-induced knockdown of SRF using Western blot (**Fig. S2**). Next, to stain surface receptors, we used the membrane-impermeant chemical crosslinking reagent *BS^3^,* which selectively distinguishes cell surface proteins from high-molecular-mass aggregates (Boudreau *et al*, 2012; Boudreau & Wolf, 2005). The immunoblot analysis of AMPARs in crosslinked neurons revealed a decrease in surface GluR1 and GluR2 subunit levels in cultures that were transduced with AAV-shSRF particles compared with AAV-shCTR (**Fig. 5B, C**; sGluR1: AAV-shCTR, 1.00 ± 0.04, *n* = 8, AAV-shSRF, 0.37 ± 0.05, *n* = 8, Student’s *t*-test, *p* < 0.0001; sGluR2: AAV-shCTR, 1.00 ± 0.068, *n* = 8; AAV-shSRF, 0.54 ± 0.13, *n* = 8, Student’s *t*-test, *p* = 0.0076). To test whether the decrease in surface GluR1 and GluR2 levels was attributable to the downregulation of total AMPAR protein levels, Western blot analysis was performed using independent extracts from the same biological experiment. Similar to the crosslinking experiments, cultured hippocampal neurons were transduced with AAV-shSRF or AAV-shCTR on DIV5, and total protein levels were analyzed on DIV21 (**Fig. 5E, F**; GluR1: AAV-shCTR, 1.00 ± 0.05, *n* = 8; AAV-shSRF, 0.32 ± 0.05, *n* = 8, Student’s *t*-test, *p* < 0.0001; GluR2: AAV-shCTR, 1.00 ± 0.07, *n* = 8; AAV-shSRF, 0.56 ± 0.12, *n* = 8, Student’s *t*-test, *p* = 0.0061). We found that the total protein level of GluR1 and GluR2 also significantly decreased in SRF-depleted neurons compared with control cells.

**Fig 5.**
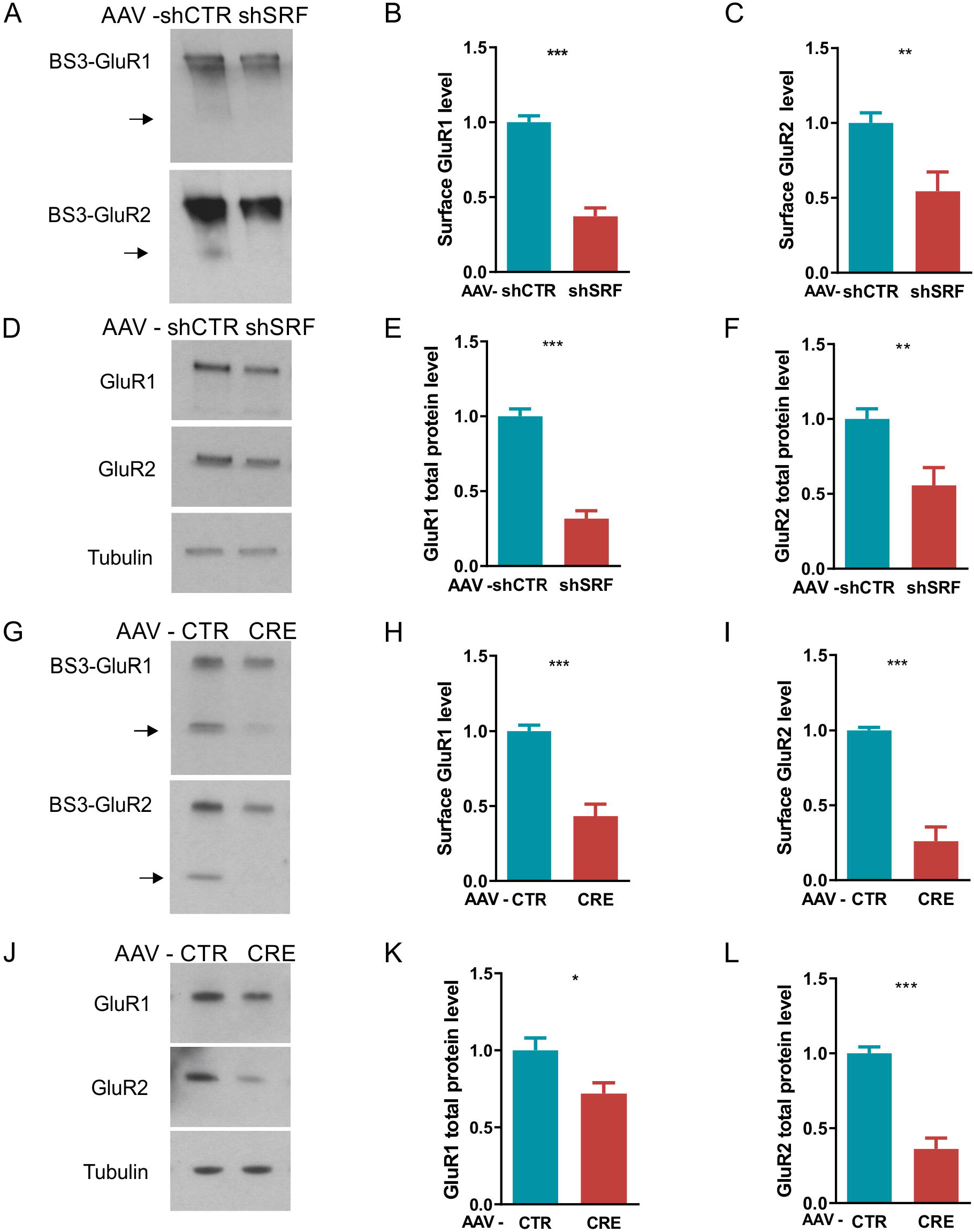
SRF controls surface and total protein levels of GluR1 and GluR2 AMPAR subunits. **(A)** Western blots of surface proteins in extracts from AAV-shCTR- and AAV-shSRF-transduced rat hippocampal neurons. Surface GluR1 and GluR2 were BS^3^ crosslinked before protein extraction. Arrows indicate the intracellular poll of receptors. **(B, C)** Quantification of surface GluR1 and GluR2 protein levels in AAV shCTR- and shSRF-transduced rat neurons. **(D)** Western blot analysis of GluR1 and GluR2 total protein expression in AAV**-** shCTR- and AAV**-** shSRF-transduced neurons. **(E, F)** Quantification of total GluR1 and GluR2 protein levels in AAV-shCTR- and AAV-shSRF-transduced rat neurons. **(G)** Western blots of BS^3^ crosslinked surface proteins in extracts from CTR- and CRE-AAV-transduced *Srf*^f/f^ hippocampal neurons. Arrows indicate the intracellular poll of receptors. **(H, I)** Quantification of surface GluR1 and GluR2 protein levels in AAV-CTR- and AAV-CRE-transduced *Srf*^f/f^ mouse neurons. **(J)** Western blot analysis of GluR1 and GluR2 total protein expression in AAV**-** CTR- and AAV-CRE-transduced mouse hippocampal neurons. **(K, L)** Quantification of total GluR1 and GluR2 protein levels in AAV-CTR- and AAV-CRE-transduced mouse neurons. Tubulin served as the loading control. *\*p* < 0.05, *\*\*p* < 0.01, *\*\*\*p* < 0.001 (Student’s *t*-test). All of the data are presented as a fold change relative to control. The data are from at least three independent neuronal cultures and are expressed as mean ± SEM.

Next, to verify the results that were obtained by shRNA, we performed an independent analysis using mouse neurons that were isolated from the SRF^f/f^ mouse hippocampus that was transduced with AAV-Cre recombinase or AAV-CTR on DIV5-6. Similar to neurons that were transduced with AAV-shRNA, SRF deletion by CRE recombinase expression decreased both membrane-bound GluR1 and GluR2 (**Fig. 5H, I**; sGluR1: AAV-CTR, 1.00 ± 0.04, *n* = 5; AAV-CRE, 0.43 ± 0.08, *n* = 5, Student’s *t*-test, *p* = 0.0002; sGluR2: AAV-CTR, 1.00 ± 0.02, *n* = 5, AAV-CRE, 0.26 ± 0.08, *n* = 5, Student’s *t*-test, *p* < 0.0001) and total protein levels (**Fig. 5K, L**; GluR1: AAV-CTR, 1.00 ± 0.08, *n* = 8; AAV-CRE, 0.72 ± 0.07, *n* = 8, *t*-test, *p* = 0.0202; GluR2: AAV-CTR, 1.0 ± 0.04, *n* = 5; AAV-CRE, 0.36 ± 0.07, *n* = 5, *t* -test, *p* < 0.0001). The surface and total protein expression of GluR1 and GluR2 subunits were significantly reduced by SRF deletion in both *in vitro* models.

### Downregulation of Gria1 and Gria2 mRNA expression in SRF-depleted neurons

The molecular mechanisms by which SRF regulates synaptic plasticity are linked to its transcriptional activity and regulation of expression of several activity-induced genes. To investigate whether SRF controls GluR1 (encoded by the *Gria1* gene) and GluR2 (encoded by the *Gria2* gene) gene expression during neuronal maturation, we conducted qPCR. We evaluated the expression of *Gria1* and *Gria2* mRNA in hippocampal neurons that were transduced with AAV-shSRF and AAV-shCTR on DIV5. Both mRNAs that encode AMPARs were downregulated in extracts from SRF-depleted rat hippocampal neurons on DIV21 (**Fig. 6A**; *Gria1*: AAV-shCTR, 0.99 ± 0.04, *n* = 8; AAV-shSRF, 0.67 ± 0.06, *n* = 8, Mann-Whitney test, *p* = 0.0006; *Gria2*: AAV-shCTR, 0.99 ± 0.03, *n* = 8; AAV-shSRF, 0.77 ± 0.04, *n* = 8, Mann-Whitney test, *p* = 0.0132). We also found that AAV-shSRF significantly downregulated *Srf* mRNA levels (**Fig. 6A**; *Srf*: AAV-shCTR, 1.02 ± 0.03, *n* = 8; AAV-shSRF 0.38 ± 0.04, *n* = 8; Mann-Whitney test, *p* < 0.0001).

**Fig. 6.**
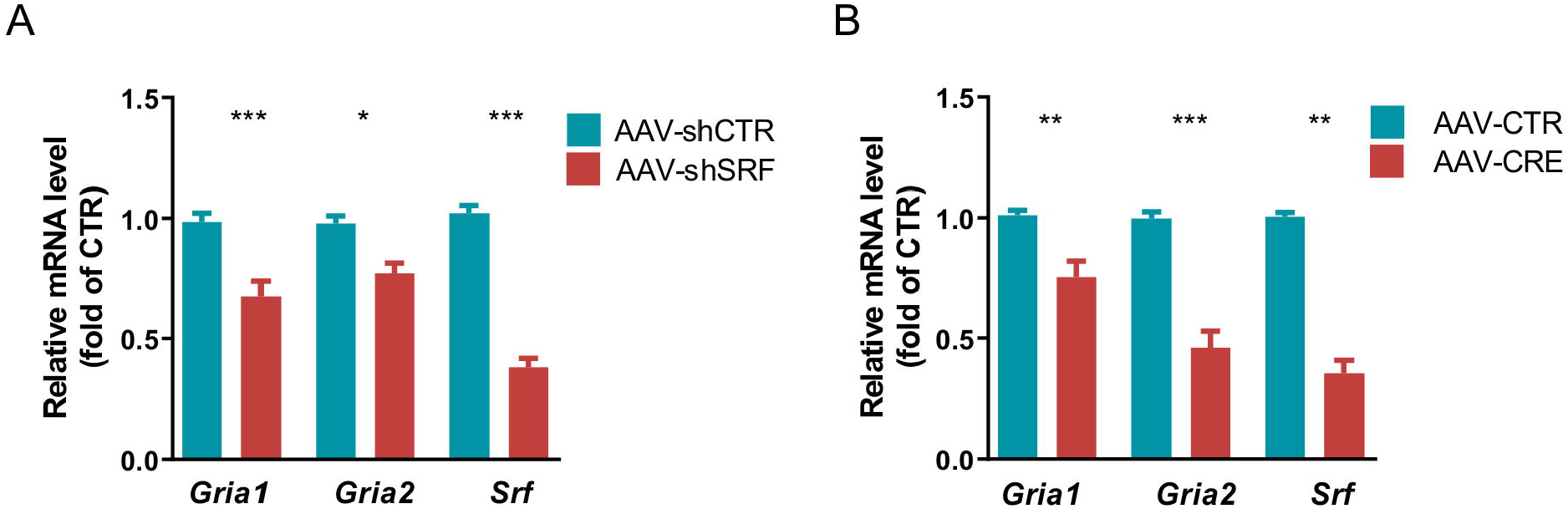
Lack of SRF reduces *Gria1* and *Gria2* expression in rat and mouse hippocampal neurons. mRNA levels were quantified in hippocampal neurons with SRF deletion during neuronal development in rat and mouse neuronal cultures. **(A)** q-RT-PCR analysis of *Gria1*, *Gria2*, and *Srf* mRNA expression in rat hippocampal neurons that were transduced with AAV-shCTR or AAV-shSRF. **(B)***Gria1*, *Gria2*, and *Srf* mRNA quantification in AAV-CTR- and AAV-CRE-transduced *Srf*^f/f^ mouse neurons. *\*p* < 0.05, *\*\*p* < 0.01, *\*\*\*p* < 0.001 (Mann-Whitney test). The data are from at least three independent rat or mouse cultures and are expressed as mean ±SEM. The *Gria2* and *Srf* data from rat cultures in **(A)** passed the normality test. *Gria2*: *\*\*p* = 0.0016; *Srf*: *\*\*\*p* < 0.0001 (Student’s *t*-test). The *Gria1*, *Gria2*, and *Srf* data from mouse cultures in (**B**) passed the normality test. *Gria 1*: *\*\*p* = 0.0037; *Gria2*: *\*\*\*p* < 0.0001; *Srf*: *\*\*\*p* < 0.0001 (Student’s *t*-test).

Moreover, the results were validated by an independent analysis using SRF^f/f^ mouse neurons from the hippocampus that were transduced with AAV-Cre recombinase particles or AAV-CTR on DIV 5-6. Similar to the shRNA results, SRF deletion in mouse neurons decreased both *Gria1* and *Gria2* mRNA levels (**Fig. 6B**; *Gria1*: AAV-CTR, 1.01 ± 0.02, *n* = 7; AAV-CRE, 0.75 ± 0.06, *n* = 8, Mann-Whitney test, *p* = 0.0011; *Gria2*: AAV-CTR, 0.99 ± 0.03, *n* = 8; AAV-CRE, 0.46 ± 0.07, *n* = 8, Mann-Whitney test, *p* = 0.0002; *Srf*: AAV-CTR, 1.00 ± 0.02, *n* = 6; AAV-CRE, 0.35 ± 0.05, *n* = 6, Mann-Whitney test, *p* = 0.0022). These results suggest that SRF regulates AMPAR expression at the transcriptional level.

### SRF controls spine maturation in the hippocampus in vivo and social behaviors

To assess the role of SRF in spine maturation *in vivo*, we used conditional SRF mutants (SRF^f/f^) that were crossed with a tamoxifen-inducible CRE recombinase line under the CaMKIIα promoter (CaMKCreER^T2^). Starting on P5-6, mouse pups were injected with one or three doses of 4-OHT, every other day, to stimulate the translocation of Cre recombinase to the nucleus and *SRF* deletion (**Fig. 7A**). Next, young adult *Srf ^f/f^* (wildtype, WT) and *Srf* gene KO mice that carried a single copy of Cre recombinase *Srf ^f/f;CaMKCreERT2^* (KO) were used for the experiments.

**Fig. 7.**
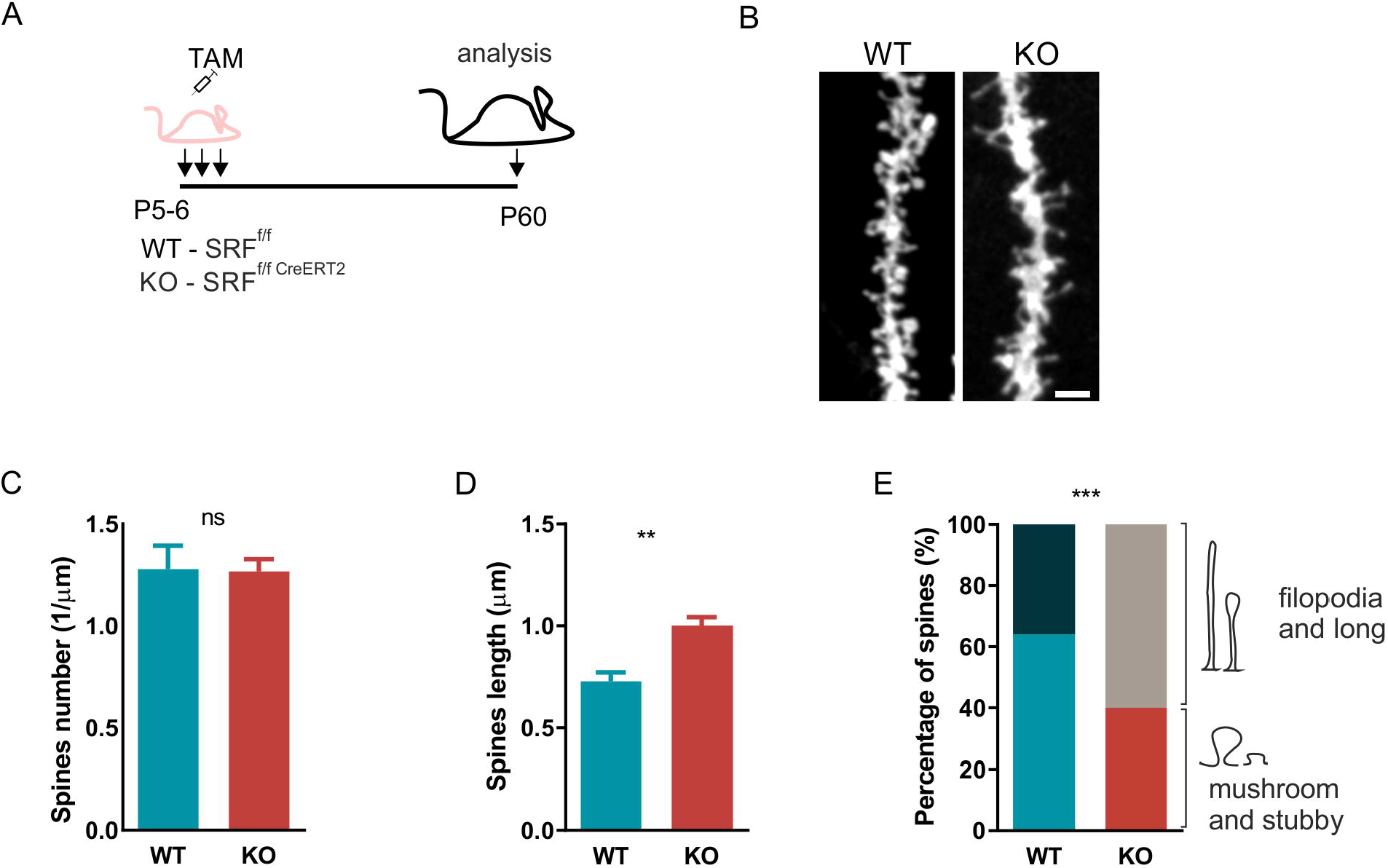
Early postnatal deletion of SRF affects spine maturation in the hippocampal CA1 neurons *in vivo*. **(A)** Schematic diagram of the experimental design of the early postnatal deletion of SRF in excitatory neurons. **(B)** Representative confocal images of DiI-stained dendrites with dendritic spines of neurons from the CA1 area in wildtype (WT) and SRF KO animals. Scale bar = 2 μm. **(C, D)** Dendritic spine density and length in neurons from WT and SRF KO animals. **(E)** Percentage of protrusions that clustered into two categories (“filopodia and long spines” and “mushroom and stubby spines”) in the hippocampal CA1 area. *^ns^p* > 0.05, *\*\*p* < 0.01, *\*\*\*p* < 0.001 (Student’s *t*-test). Data from *n* = 4 WT mice and *n* = 7 KO mice are shown. The data are expressed as mean ± SEM.

To examine the consequences of developmental deletions of SRF in hippocampal excitatory neurons, we measured the density and morphology of dendritic spines in the stratum radiatum of the hippocampal CA1 region in adult animals (**Fig. 7B**). The analysis of spine density showed no significant changes in the number of spines (**Fig. 7C**; WT: 1.28 ± 0.11, *n* = 4; KO: 1.27 ± 0.06, *n* = 7; Student’s *t*-test, *p* = 0.0248). However, we observed a substantial increase in dendritic spine length in SRF KO animals (**Fig. 7D**; WT: 0.72 ± 0.04, *n* = 4; KO: 1.00 ± 0.04, *n* = 7; Student’s *t*-test, *p* = 0.0021). To investigate how SRF deletion affects dendritic spine shape, we clustered spines into two morphological categories (“filopodia and long spines” and “mushroom and stubby”) as previously done *in vitro* (see **Fig. 1F, G**). We found that SRF KO mice exhibited a significant 24% increase in the frequency of filopodia and long spines and a decrease in the population of mushroom spines in the CA1 (χ^*2*^ = 11.54, *df* = 1, *p* = 0.0007, *χ*^2^ test, *p* = 0.0007) compared with wildtype animals.

Alterations of spine morphology and changes in the expression of glutamate receptors are frequently linked to neurodevelopmental disorders, among which ASD is the most prevalent (Penzes *et al.*, 2011; Phillips & Pozzo-Miller, 2015; Soto *et al*, 2014). Thus, considering the presence of structural changes in dendritic spines in SRF KO animals, we hypothesized that an early postnatal lack of SRF affects some aspects of social behavior, impairments of which are core symptoms of the autistic phenotype that is frequently observed in animal studies of the disorder (Silverman *et al*, 2010). We used Eco-HAB, an RFID-based system for the automated assessment of individual, voluntary social behavior in group-housed mice under semi-natural conditions (**Fig. 8A**). Eco-HAB was previously shown to accurately evaluate social deficits in mice (Puscian *et al*, 2016). Cohorts of male WT and SRF KO mice were subjected to 96 h of testing, and behavior was continuously recorded (**Fig. 8B**). Social behavior was measured after 24h of habituation to the novel Eco-HAB environment when the group’s social structure stabilized. We found that SRF KO animals exhibited severe impairments in the pattern of interactions with familiar mice (in-cohort sociability), measured as the time voluntarily spent together with other animals within the group (**Fig. 8C-E**). SRF KO mice were less willing to spend time together compared with WT animals (**Fig. 8 C**; WT 0.08 ± 0.006; SRF KO: 0.01 ± 0.002, *n* = 11, Mann-Whitney test, *p* < 0.0001; **Fig. 8 D**; Kolmogorov-Smirnov test, *p* < 0.0001). Thus, male SRF KO mice modeled a specific impairment that is found in the majority of autistic patients who are unable to establish and maintain social contacts (Chasson G., 2014). Despite this, interest in novel social stimuli, measured as a proportion of time spent exploring social *vs*. non-social scents that were presented behind the perforated partitions of the opposing Eco-HAB cages, was robust and had a similar intensity in WT and SRF KO male mice (**Fig. 8F**; WT: 2.36 ± 0.5, *n* = 11; SRF KO: 2.83 ± 0.39, *n* = 11; Mann-Whitney test, *p* = 0.1932).

**Fig. 8.**
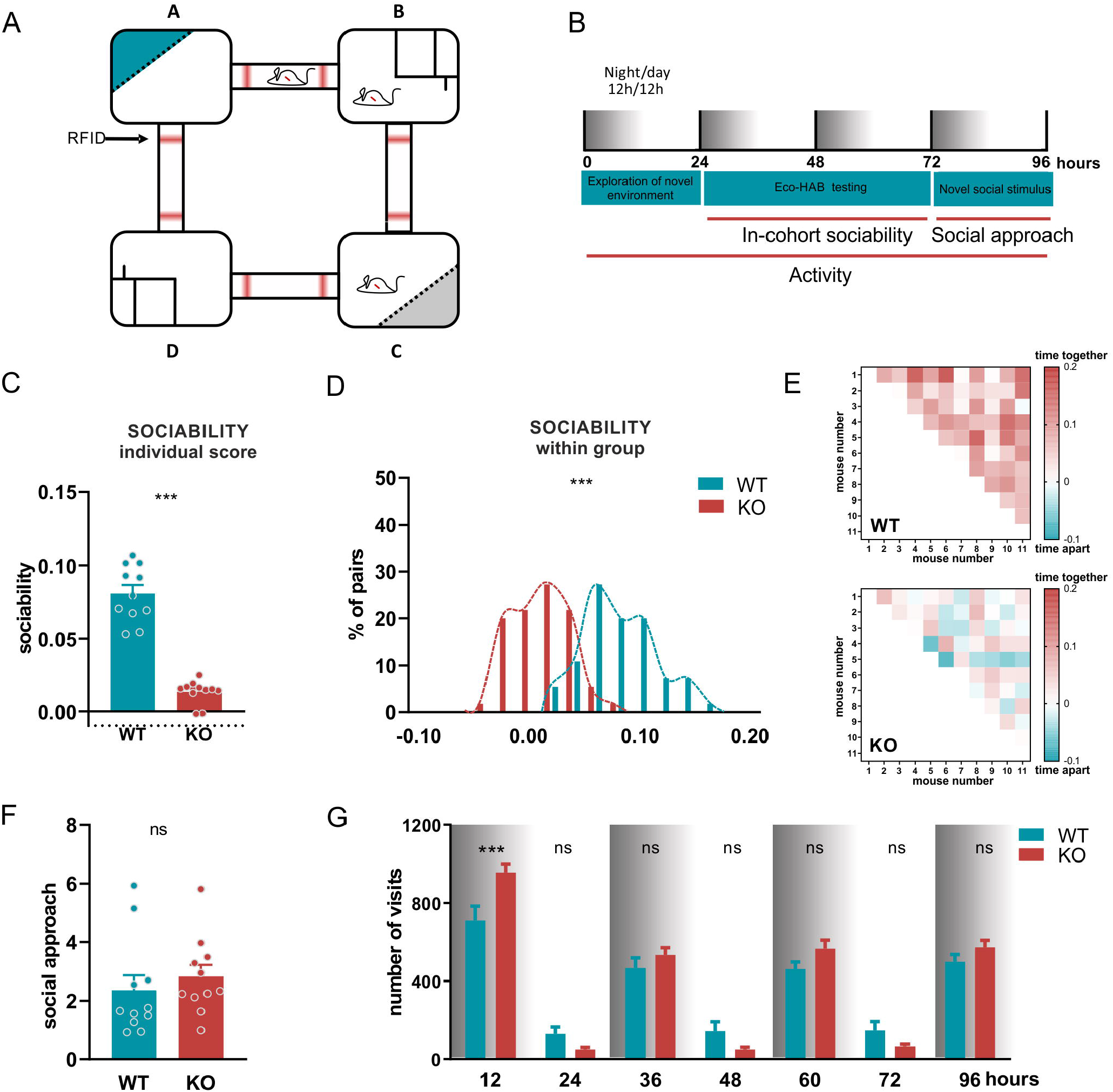
SRF KO mice exhibit decrease in sociability despite normal response to novel social stimuli. **(A)** Animals were tested in Eco-HAB, an RFID-based system for the automated assessment of voluntary social behavior in group-housed mice under semi-naturalistic conditions. The system consists of four housing compartments that are connected by tube-shaped passages that are equipped with RFID antennas that track individual behavior of each mouse. Compartments B and D contain food and water, and compartments A and C contain impassable, perforated partitions (dashed lines), behind which social (blue) and non-social (gray) cues are presented. **(B)** Experimental timeline. Mice were housed in the Eco-HAB chamber for 96 h and tested for exploration of the novel environment, in-cohort sociability, approach to novel social odor, and general locomotor activity. **(C)** Sociability, measured as the time the mouse voluntarily spent together in WT and SRF KO animals. **(D)** Histogram of the distribution of in-cohort sociability in WT and SRF KO mice. \*\*\**p* < 0.0001 ( Kolmogorov-Smirnov test). **(E)** Raw data from **(C)** and **(D)**, represented as a matrix in which squares illustrate the time voluntarily spent together by each pair of mice within the cohort. The intensity of colors reflects the strength of the relationship in accordance with the presented scale. **(F)** Social approach in WT and SRF KO mice. *^ns^p* > 0.05, *\*\*\*p* < 0.001 in **(C)** and **(F)**(Mann-Whitney test). The data are expressed as mean ± SEM. (**G**) Activity in WT and SRF KO mice in the dark and light phases during 96 h of Eco-HAB recording. Two-way ANOVA with post-hoc Sidak’s multiple comparisons test; ***p < 0.0001; n = 11 male WT mice; n = 11 male SRF KO mice. The data are expressed as mean ± SEM.

Notably, no differences in overall activity were observed between WT and SRF KO animals when social behavior was assessed. Thus, the reported differences were not attributable to that factor (**Fig. 8G**; two-way repeated-measures ANOVA: genotype: *F*_1,10_ = 3.98, *p* = 0.074 genotype, time: *F*_6,60_ = 229.1, *p* < 0.0001; genotype × time: *F*_6,60_ = 10.76, *p* < 0.0001; Sidak multiple-comparison *post hoc* test, ^ns^*p* > 0.05, \*\*\**p* < 0.001). However, consistent with previous results (Nader *et al.*, 2019), we found that SFR KO animals exhibited a hyperactive pattern of exploration of the novel environment, reflected by an increase in the number of visits to all Eco-HAB compartments in the first 12 h of the experiment (**Fig. 8G**).

As currently recommended, we tested the SRF KO phenotype in both sexes (Beery, 2018; Clayton, 2016). We applied the previously described approach to evaluate social behavior in SRF KO females. SFR KO females exhibited changes in both measures, including a decrease in in-cohort sociability (**Fig. S3A-C**; WT: 0.05 ± 0.004, *n* = 12; SRF KO: 0.03 ± 0.002, *n* = 11; Mann-Whitney test, *p* = 0.0028; Kolmogorov-Smirnov test, *p* < 0.0001) and an increase in interest in novel social stimuli (**Fig. S3D**; WT: 1.62 ± 0.31, *n* = 12; SRF KO: 2.61 ± 0.35, *n* = 11; Mann-Whitney test, *p* = 0.0317). These results further confirmed the specificity of the observed social deficits. Although female SRF KO mice exhibited an increase in social approach **(Fig. S3D**), they did not maintain a normal pattern of interactions with familiar co-housed mice, reflected by the decrease in time voluntarily spent together with others. These data confirm the need to test phenotypes of both sexes and are consistent with previous findings that showed different symptoms in autistic females (Dworzynski *et al*, 2012; Mandy *et al*, 2012; Ratto *et al*, 2018; Werling & Geschwind, 2013).

## Discussion

In this study, we present a coherent picture of hippocampal excitatory synapse function at the morphological, biochemical, and physiological levels. We found that SRF expression during the postnatal development of hippocampal neurons is necessary for proper synapse maturation, which has consequences at the behavioral level.

Using two *in vitro* models and one *in vivo* model, we found that the early postnatal deletion of SRF in excitatory neurons impaired the morphological and functional maturation of dendritic spines. SRF knockdown in developing hippocampal excitatory neurons resulted in an increase in the length of spines and number of immature filopodia-like protrusions, with a general lack of changes in the overall density of dendritic spines *in vitro* and *in vivo*. These changes in spine shape might be related to the decrease in β-actin levels and cofilin-1 inactivation upon SRF ablation, which was previously reported by us and others (Alberti *et al.*, 2005; Beck *et al*, 2012; Nader *et al.*, 2019; Zimprich *et al*, 2017). Notably, both proteins play a fundamental role in dendritic spine expansion (Bosch *et al*, 2014; Zimprich *et al.*, 2017). The alterations of spine morphology appear to directly translate into a reduction of the number of functional synapses, as we observed a decrease in the synaptic colocalization of well-known markers of excitatory synapses, Bassoon and PSD-95, in SRF-depleted neurons. Moreover, the reduction of SRF expression significantly decreased the number of PSD-95 puncta. PSD-95 is a major postsynaptic scaffold protein that modulates postsynaptic function and the maturation of excitatory synapses (Ehrlich *et al*, 2007; El-Husseini *et al*, 2000; Taft & Turrigiano, 2014). Appropriate levels of PSD-95 expression were shown to be required for structural and functional synaptic development (Ehrlich et al., 2007). Altogether, our results suggest that thin filopodia-like protrusions in SRF-depleted neurons create fewer functional synapses, with no changes in overall spine density. This conclusion, together with the reduction of the frequency of AMPAR-mediated mEPSCs that were recorded from SRF-depleted neurons, may indicate the presence of silent synapses (i.e., immature synapses that do not possess functional AMPARs; (Isaac *et al*, 1995; Liao *et al*, 1995). Such structures are highly abundant in juvenile brain networks and disappear during development either by acquiring AMPARs or by being pruned from the dendrite. The reduction of mEPSC frequencies, with no changes in dendritic spine density, suggests a role for SRF in the maturation of dendritic spines into functional synaptic contacts.

The mechanisms that drive the postnatal functional maturation of neurons are poorly understood, although the expression of neurotransmitter receptors and composition of their subunits have been linked to neuronal maturation (Lohmann & Kessels, 2014; Monyer *et al*, 1991; Pickard *et al*, 2000; Sheng *et al*, 1994; Williams *et al*, 1993). Interestingly, genes that are upregulated in late postnatal stages of hippocampal development correspond to the peak of synapse maturation, including such glutamate receptors as *N*-methyl-D-aspartate receptors (NMDARs) GluR1, and GluR2 (Mody *et al*, 2001), the time- and region-specific expression of which significantly influences dendritic morphology and synaptic connectivity (Chen *et al*, 2009). We found that SRF, a primary regulator of activity-induced gene expression (Kuzniewska *et al.*, 2016; Kuzniewska *et al.*, 2013; Parkitna *et al.*, 2010; Ramanan *et al.*, 2005), was active during a specific time-window of postnatal brain development and essential for the expression of GluR1 and GluR2 in excitatory neurons. The number and composition of AMPARs at synapses determine their functional maturation and strength, which are fundamental mechanisms that underlie synaptic plasticity (Henley & Wilkinson, 2016; Matsuzaki *et al*, 2001; Nimchinsky *et al.*, 2002; Tracy *et al*, 2011). Moreover, the expression of AMPARs is developmentally regulated (Jourdi *et al*, 2003; Orlandi *et al*, 2011; Robertson *et al*, 2009). Our data clearly suggest that SRF is a key transcription factor that regulates dendritic spine maturation. The levels of *Gria1* and *Gria2* mRNA and surface AMPARs decreased in neurons that lacked SRF. This conclusion was additionally supported by the observation of a lower amplitude of mEPSCs in SRF-depleted neurons. Our results are consistent with data that showed that the loss or overexpression of postsynaptic AMPARs altered overall synaptic transmission in developing neurons and affected synapse morphogenesis and maturation (Passafaro *et al*, 2003; Saglietti *et al*, 2007; Tracy *et al.*, 2011). Moreover, a reduction of PSD-95-positive synaptic puncta in SRF-depleted neurons could contribute to the smaller number of AMPARs at synapses. PSD-95 is essential for their synaptic accumulation and AMPAR-mediated responses (Buonarati *et al*, 2019; Chen *et al*, 2015; Nair *et al*, 2013). Interestingly, mutations and deletions of genes that encode AMPAR subunits, mainly GluR2, were recently linked to such neuropsychiatric conditions as ASD and mental disabilities (Hackmann *et al*, 2013; Ramanathan *et al*, 2004; Salpietro *et al*, 2019; Soto *et al.*, 2014). Altogether, our data suggest that SRF plays a critical role in converting filopodia to functional synapses by controlling the level of AMPARs. Remaining to be investigated is whether SRF is directly or indirectly involved in the regulation of *Gria1* and *Gria2* gene transcription in neurons. Interestingly, SRF was recently shown to bind *Gria1* and *Gria2* promotor regions in hippocampal neurons (Del Blanco *et al.*, 2019). We do not exclude the possible impact of other neuronal SRF targets on the regulation of AMPAR expression (Kim *et al*, 2018; Knoll & Nordheim, 2009). Our previous results demonstrated that SRF regulates the expression of c-*fos*, a component of the AP-1 transcription factor complex (Kuzniewska *et al.*, 2016). Interestingly, activity-regulated AP1 is a crucial mediator of postnatal neuronal development. During maturation, AP-1 expression increases and correlates with the formation of new gene enhancers that increase the expression of neuronal maturation genes (Stroud *et al*, 2020). The morphological, biochemical, and physiological synaptic immaturity that was observed in our study may be a direct consequence of a decrease in the chromatin occupancy of SRF or an indirect consequence of SRF-regulated c-*fos* expression in postnatal neurons.

Our results are especially interesting in the context of a recent study by Del Blanco et al. (2019), who investigated the role of cyclic adenosine monophosphate response element binding protein (CREB)-binding protein (CBP) in spine maturation (Del Blanco *et al.*, 2019). The lack of CBP during neuronal development alters the expression of genes that are involved in neuronal growth and plasticity. Not only, many of these genes have SRF binding elements, but also deficits in dendritic spine maturation in CBP-lacking neurons were reversed by the overexpression of SRF transcription through an SRF-VP16 construct (Del Blanco *et al.*, 2019). SRF acts upstream of CBP-controlled transcription, suggesting that it is critical for dendritic spine maturation and activity-dependent synaptic remodeling (Del Blanco *et al.*, 2019). Notably, CBP is known to interact not only with CREB but also with other transcription factors that can recruit CBP to gene promoters. Several lines of evidence suggest that SRF and CBP may co-regulate gene expression (Hanna *et al*, 2009; Nissen *et al*, 2001; Qiu & Li, 2002; Ramos *et al*, 2010). Moreover, the lack of CBP in neurons decreased the expression of AMPAR and NMDAR subunits (Chen *et al*, 2010). The role of SRF in spine maturation is further supported by the function of myocardin-related transcription factor A (MRTF-A) and MRTF-B, major transcriptional co-activators of SRF (Posern & Treisman, 2006), during synapse development. The inhibition of both MRTF-A and MRTF-B significantly decreased the percentage of mature dendritic spines and increased the percentage of long immature spines (Kaneda *et al*, 2018), thus resembling the phenotype that is observed in neurons that developmentally lack SRF.

The present study focused on the hippocampus, which is not only involved in spatial memory formation, but of particular interest when studying different aspects of social interactions. Hippocampal neurons project to different brain areas that are critically engaged in the regulation of social behaviors (Phillips *et al*, 2019). The hippocampus itself can directly influence social interactions (Hitti & Siegelbaum, 2014). Moreover, hippocampus-dependent behavioral deficits in sociability are often observed in several neurodevelopmental disorders, including ASD (Kwon *et al*, 2006), and these hippocampal dendritic spines have a morphology that is similar to SRF-lacking neurons.

Overall, using a model of SRF deletion in early postnatal stages of hippocampal excitatory neuron development, we demonstrated that the SRF-dependent maturation of dendritic spines influences social behavior. To study the behavioral phenotype, we used a fully automated system, Eco-HAB, that allowed us to assess animal behavior without human presence (Puscian *et al.*, 2016). We found that both male and female SRF KO mice exhibited a specific deficit of interacting with well-known animals, in which they were less willing to spend time together with their co-housed conspecifics. SRF KO mice of both sexes were unable to establish and maintain social contacts, thus strengthening the relevance of the observed behavioral impairments. Interestingly, similar deficiencies in in-cohort measures of sociability were identified in *Fmr1* KO mice (Puscian *et al.*, 2016), a well-known model of autism (Bernardet & Crusio, 2006; Mineur *et al*, 2006; Santos *et al*, 2014). Similar to SRF KO animals, changes in the morphology of dendritic spines, indicating general spine immaturity, were also shown in *Fmr1* KO mice and patients with fragile X syndrome (Comery *et al*, 1997; Irwin *et al*, 2000; Nimchinsky *et al.*, 2001). Moreover, abnormal levels of AMPAR density and disruptions of excitatory transmission in SRF-depleted neurons might be partially responsible for the observed social phenotype. Changes in AMPAR-dependent transmission are involved in the pathophysiological mechanism of ASD, and AMPAR-related excitatory transmission can regulate social behaviors (Kim *et al*, 2019). Additionally, lower AMPAR density was associated with ASD in humans (Purcell *et al*, 2001). We recently showed that SRF deletion in adult neurons impaired hippocampus-dependent measures of social behavior, such as nest building and marble burying (Nader *et al.*, 2019). Such a behavioral phenotype in adult animals is consistent with observations of young SRF KO mice. Notably, our results showed no changes in social approach in male SRF KO mice, in contrast to female mice, in which we observed a slightly higher response to the novel olfactory cue. Sex-dependent behavioral disparities are often observed in developmental disorders, such as autism (Mandy *et al.*, 2012), and the underlying causes require further investigation. Altogether, our molecular and behavioral data suggest that changes in SRF expression might be partially responsible for some symptoms that are observed in ASD.

The present study used both *in vitro* and *in vivo* models we showed that postnatal SRF expression in excitatory neurons is necessary for proper dendritic spine maturation and neuronal properties of AMPAR-dependent transmission. Our data shed light on the transcriptional mechanism that underlies the functional maturation of neurons. We found that SRF-regulated transcription in the nucleus is crucial for postnatal neuronal maturation. Based on these data, we hypothesize that the proper time and level of transcription of genes that encode AMPARs is the first stage of synaptic maturation. Our data suggest that lower levels of AMPAR mRNAs cause reductions of the number of synaptic AMPARs and AMPAR-mediated excitatory transmission and inhibition of the morphological maturation of dendritic spines. Additionally, structural changes in dendritic spines and modifications of cell function that were observed in SRF-depleted neurons were accompanied by distinct social deficits that are commonly observed in ASD. The identification of novel molecular targets that are involved in the regulation of distinct aspects of sociability may reveal the neuronal mechanisms that underlie some neuropsychiatric diseases, such as ASD.

## Methods

### Animals

Srf KO mice (mutant mice; *Srf*^f/f*CaMKCreERT2*^) and control CreERT2-negative littermates (control mice *Srf*^f/f^) were used (Kuzniewska *et al.*, 2016; Nader *et al.*, 2019). On P5-6, *Srf*^f/f^ and *Srf*^f/f*CaMKCreERT2*^ pups were intraperitoneally injected every other day with three doses of 0.25 mg 4-hydroxytamoxifen (4-OHT; Sigma, #7904) or one dose of 0.75 mg of this compound in sunflower oil. Young adult animals (2-3 months, males and females) were used. The mice were housed in individual cages under a 12 h/12 h light/dark cycle with food and water available *ad libitum*. The studies were performed in accordance with the European Communities Council Directive of November 24, 1986 (86/609/EEC), Animal Protection Act of Poland and approved by the 1st Local Ethics Committee in Warsaw (permission no 622/2018, 951/2019 and 984/2020). All efforts were made to minimize the number of animals used and their suffering.

### Primary neuronal cell cultures

Dissociated primary hippocampal cultures were prepared from either P0 Wistar rats or P0 *Srf* ^f/f^ transgenic mice as described previously (Magnowska *et al*, 2016). Dissociated hippocampal neurons were plated at a density of 120,000 cells per 18-mm-diameter coverslip that was coated with 1 mg/ml poly-D-lysine (Sigma, #A-003-E) and 2.5 μg/ml laminin (Roche, #11243217001). The cells were cultured in a medium that contained Neurobasal A, 2% B27 supplement, Gluta-Max, 100 U/ml penicillin, and 0.1 mg/ml streptomycin at 37°C in a 5% CO_2_ atmosphere. The day of plating was considered day 0 *in vitro* (DIV0). On DIV3, cytosine arabinoside (Ara C; 2.5 μM) was added to inhibit non-neural cell growth.

### DNA constructs and transfection

The pRNAT-H1.1/Shuttle-shSRF and pRNAT-H1.1/Shuttle-shCTR plasmids, both of which drive green fluorescent protein (GFP) expression, were transfected using Lipofectamine 2000 (Invitrogen, #11668-019) on DIV6-7 according to the manufacturer’s protocol. The plasmids were provided by Dr. B. Paul Herring (Department of Cellular and Integrative Physiology, Indiana University School of Medicine, Indianapolis, Indiana, USA). For mouse cultures, CaMKIIα-GFP and CaMKIIα-Cre plasmids were transfected using Lipofectamine 2000. Dendritic spine morphology analyses were performed on DIV18-21. In the biochemical experiments that involved the transduction of adeno-associated virus (AAV1/2, isotype 1 and 2; AAV-shCTR, AAV-shSRF, AAV-CaMKIIα-mCherry, AAV-CaMKIIα-Cre), hippocampal neurons were transduced on DIV5-6 and collected for protein or RNA analysis on DIV18-21. The electroporation of freshly dissociated newborn hippocampal neurons was conducted using rat neuron nucleofection reagents (Amaxa, Lonza, Germany) with the 5×SRE-Luc reporter plasmid (Stratagene) and EF1αLacZ (β-galactosidase [β-Gal]), both of which were described previously (Kalita *et al*, 2006).

### Luciferase and β-galactosidase reporter gene assays

Luciferase and β-Gal activity were evaluated in neuronal cell lysates using commercial assay kits (Promega, #E2000, #1500). Luminescence recordings were performed using an Infinite M200 microplate reader with an injector system (Tecan). To measure β-Gal activity, absorbance at 420 nm was read with a Sunrise 96-well plate reader (Tecan). Transcriptional activity was determined as luciferase activity normalized to β-Gal activity and compared with SRF-driven transcription on DIV3.

### Immunostaining and confocal microscopy

Hippocampal cultures were fixed for 8 min at room temperature with 4% paraformaldehyde and 4% sucrose in phosphate-buffered saline (PBS), washed with PBS, permeabilized for 10 min with 0.1% Triton X-100 in PBS, and blocked for 2 h at room temperature with 10% normal goat serum in PBS. After blocking, the cells were incubated overnight with mouse anti-GFP antibody (Millipore, #MAB3580), anti-Bassoon antibody (SySy, #141 003), anti-PSD-95 antibody (Millipore, #MABN68), or anti-SRF antibody (Santa Cruz, # sc-13029) at 4°C. The cells were then washed in PBS and incubated with fluorescent Alexa Fluor-488-, Alexa Fluor-546-, or Alexa Fluor-555-conjugated secondary antibody (Invitrogen, #A 11008; #A 21202, #A 10040, #A 32727) for 1 h at room temperature and washed and mounted with Fluoromount-G (SouthernBiotech, #0100-01). For dendritic spine analysis, GFP-positive pyramidal neurons were examined under a confocal microscope (Leica TCS SP8) that was equipped with an HC PL APO CS2 63×/1.40 oil immersion objective using the 488-nm line of an argon laser. The pixel resolution was 2048×2048, and the resulting pixel size was 0.07 μm. The sum of *Z*-stacks (maximum intensity projections) of secondary dendrites was analyzed. For the analysis of synaptic markers, GFP-positive pyramidal neurons were acquired on a Zeiss LSM 800 Airyscan confocal microscope with a PL APO 63×/1.4 oil immersion objective using 488/561/640 nm diode lasers with sequential acquisition settings of 1024×1024 resolution, 2× optical zoom, and 0.05×0.05 μm pixel size. The settings were kept the same for all scans.

### Dendritic spine morphology analysis

Confocal images of selected dendritic spine segments were semiautomatically analyzed using SpineMagick! software (Ruszczycki *et al*, 2012). To minimize possible systematic differences, the analysis was performed on dendritic spines that belonged to secondary dendrites. The density of spines was calculated as the number of spines per 1 μm of dendrite length. Spine shapes were divided into clusters and then sorted into two groups, “filopodia and long” and “mushroom and stubby,” using custom scripts (Jasinska *et al*, 2016).

### Surface receptor staining of living hippocampal neurons

Hippocampal cultures from P0 rats were transfected with pRNAT-H1.1/Shuttle-shSRF or pRNAT-H1.1/Shuttle-shCTR plasmid as described above. Levels of GluR1- or GluR2-containing AMPARs were assessed on DIV18. To label surface AMPARs, anti-GluR1 or anti-GluR2 antibody (Sigma, #ABN241 and #MAB397, respectively), diluted to a final concentration of 10 μg/ml, were added to the neuronal cultures and incubated at 37°C for 15 min. Unbound antibodies were quickly washed with ice-cold Minimum Essential Medium MEM) and then the cells were placed in fixation medium (4% formaldehyde/4% sucrose/phosphate buffer [PB]) for 7 min. Alexa Fluor-555 or Alexa Fluor-546-conjugated secondary antibodies (1:100; Invitrogen, #A 10040, #A 32727) for appropriate species were diluted in GDB buffer (0.1% bovine serum albumin, 17 mM PB, and 0.45 M NaCl) without Triton and incubated at room temperature for 1 h. After washing, coverslips were mounted with ProLong Gold (Invitrogen, #P10144). Transfected, GFP-positive pyramidal neurons were examined under a Leica TCS SP8 confocal microscope according to the previous experiments.

### Image analysis of synaptic markers and surface receptor staining

*Z*-stacks (Maximum Intensity projections, 16 bit) of secondary and tertiary dendrites of GFP-positive neurons were processed using ImageJ software (National Institutes of Health). Based on the GFP signal, the mask of transfected neurons was created and applied to other fluorescent channels. For each neuron, at least three dendritic segments were selected for further analysis. The signal from the masked segments of each channel was thresholded based on a subjective evaluation of real puncta “clusters” compared with background noise. The same threshold was kept for each experiment. To separate overlapping objects, the “*Watershed separation*” function was implemented, and then the “*Analyze Particles*” option was applied. The pixel area size was set based on experimenter picture evaluation to exclude anything that was not an object of interest in the image. The results were then added to ImageJ “*Manager*” and then restored over each fluorescent channel to analyze the area, fluorescence intensity, and density of particles. The average value of each parameter was created per one analyzed cell.

To analyze the colocalization of Bassoon/PSD-95 within dendritic spines, the Pearson correlation coefficient was used. The analysis was performed using the SpineMagick! program. The value of the coefficient varies from −1 to 1. Values closer to 1 indicate a higher degree of colocalization. Values closer to 0 represent a lack of correlation between the signals.

### Electrophysiology

The patch-clamp technique was used to measure miniature excitatory postsynaptic currents (mEPSCs). Hippocampal neurons were grown on glass coverslips and incubated in artificial cerebrospinal fluid solution (ACSF; 119 mM NaCl, 2.5 mM KCl, 1.3 mM MgCl_2_, 1 mM NaH_2_PO_4_, 26 mM NaHCO_3_, 20 mM D-glucose, and 2.5 mM CaCl_2_, saturated with carbogen) supplemented with 100 μM picrotoxin (Abcam, #U 93631) and 0.5 μM tetrodotoxin (Tocris, # 078) and heated to 31°C. GFP-positive pyramidal cells were identified visually and patched with borosilicate glass capillaries (4-6 MΩ resistance) that were filled with Cs-based internal solution (130 mM Cs-gluconate, 20 mM HEPES, 3 mM TEA-Cl, 0.4 mM EGTA, 4 mM Na_2_ATP, 0.3 mM NaGTP, and 4 mM QX-314Cl, pH 7.0-7.1; osmolarity: 290-295 mOsm). To measure mEPSCs, approximately 20-min-long voltage-clamp recordings were collected. Miniature events were analyzed using MiniAnalysis software (Synaptosoft) with an amplitude detection threshold set to 7 pA. All mini-events that were automatically detected by the software were visually verified by the experimenter.

### BS^3^ staining

To assess surface protein levels, the crosslinking with bis(sulfosuccinimidyl)suberate (BS^3^; ThermoFisher, # 21580) protocol was applied according to (Boudreau *et al.*, 2012). Briefly, AAV-infected cultures, after three washes with fresh maintenance medium, were incubated with 2 mM BS^3^ for 10 min with agitation at 37°C. Crosslinking was terminated by quenching the reaction with 100 mM glycine (10 min, 4°C). Protein extracts were then probed using standard Western blot procedures.

### Western blot analysis

Twenty micrograms of protein extracts were run on polyacrylamide gels (BioRad, #4569033) under reducing conditions (Nader *et al.*, 2019). The standard Western blot procedure was performed using anti-GluR1 (Sigma, #ABN241), anti-GluR2 (Sigma, #MAB397), and anti-SRF (Santa Cruz, #sc-13029) antibodies. To monitor equal total protein levels, the blots were re-probed with α-tubulin (Sigma, # T9026) antibodies. For signal detection, the chemiluminescent method was used. To quantify individual bands, a scan of X-ray films was analyzed by densitometry using GeneTools software (Syngene).

### RNA preparation and quantitative real-time polymerase chain reaction

Total RNA was isolated from rat or mouse hippocampal cultures using the RNeasy Mini Kit (Qiagen, #74104). DNA contamination was removed by digestion with DNase I (Qiagen, #1023460). The RNA concentration was analyzed using NanoDrop (ThermoFisher). RNA was reverse transcribed with SuperScript IV Reverse Transcriptase (Invitrogen #18090050) according to the manufacturer’s instructions. cDNA was amplified with TaqMan probes (ThermoFisher) that were specific for mouse or rat. Quantitative real-time polymerase chain reaction (PCR) was performed using Fast TaqMan Master Mix (Applied Biosystems, #44456) with an Applied Biosystems 7900HT Fast Real-Time PCR System. Fold changes in expression were determined using the ΔΔCT relative quantification method. The values were normalized to relative amounts of Gapdh.

### Eco-HAB testing of social behaviors

To test social behavior in *Srf* KO mice, we used Eco-HAB (Puscian *et al.*, 2016), a radio frequency identification (RFID)-based system that is fully automated and allows the efficient evaluation of animals’ social phenotypes with no contact between tested animals and experimenters. The group-housed animals (SRF KO and littermate controls, 2-3 months old) were housed under a 12 h/12 h light/dark cycle with unlimited access to food and water. Mice were subjected to the 96-h testing protocol, which consisted of an adaptation phase (72 h) and the subsequent presentation of a novel social stimulus (24 h). During the latter, a novel social odor (i.e., bedding soaked in a scent of an unfamiliar mouse of the same strain, sex, and age) and neutral olfactory cue (i.e., clean bedding) were put behind the perforated partitions of the opposing Eco-HAB chambers for the animals to voluntarily explore. Data were continuously recorded throughout the experiment and then evaluated. We analyzed activity, defined as the number of visits to all four Eco-HAB compartments, calculated in 12-h time bins, relative to the alternating dark and light phases of the light/dark cycle. To assess the exploration of a novel environment, we evaluated the animals’ activity during the first 12 h upon introduction of the cohort to the Eco-HAB chambers. We also analyzed in-cohort sociability, reflecting the tendency to voluntarily spend time with conspecifics, during the second and third dark phase of adaptation (i.e., during the period when the social structure of the tested cohort stabilized). As previously described by (Puscian *et al.*, 2016), in-cohort sociability was calculated as the total time that each pair of mice spent together minus the time that the animals would spent jointly because of their individual preferences for occupying certain spaces within the Eco-HAB chamber. We also evaluated approach to social odor, which was calculated as an increase in the proportion of time that each animal spent in the compartment with a novel social odor to the time spent in the compartment with a neutral-olfactory cue relative to the respective period from the preceding adaptation phase. For detailed information about the analyses of these measures, see Puscian et al. (2016).

### Statistical analyses

To compare the distributions of datasets, the Shapiro-Wilk normality test was performed. Unpaired Student’s *t*-test or the Mann-Whitney test (nonparametric) was used to test differences between two groups. When required, one- or two-way analysis of variance (ANOVA) was performed, followed by Sidak’s multiple-comparison *post hoc* test. To compare cumulative distributions of data, the Kolmogorov-Smirnov test was used. The number of neurons and independent cultures or animals that were used for the analyses are reported in the Results section. Values of *p* < 0.05 were considered statistically significant. The results were analyzed using GraphPad Prism software.

## Acknowledgments

This work was supported by the National Science Centre, Poland (grants no. SONATA BIS 2 2012/07/E/NZ3/01814 and Opus 2019/33/B/NZ4/01450).

## Author contributions

A.K., M.R., L.M., A.B., P.M., K.N., M.P., J.J., Lu.K., A.P., and K.K. performed the experiments and analyzed the data. M.R., A.K., A.P., E.K., and KK wrote the manuscript. L.K., A.P., and E.K. discussed the data. K.K. designed the experiments, provided funding and overall supervision of this project.

## Conflict of interest

The authors report neither financial interests nor conflicts of interest.

## Figure legend

**Fig. S1.**
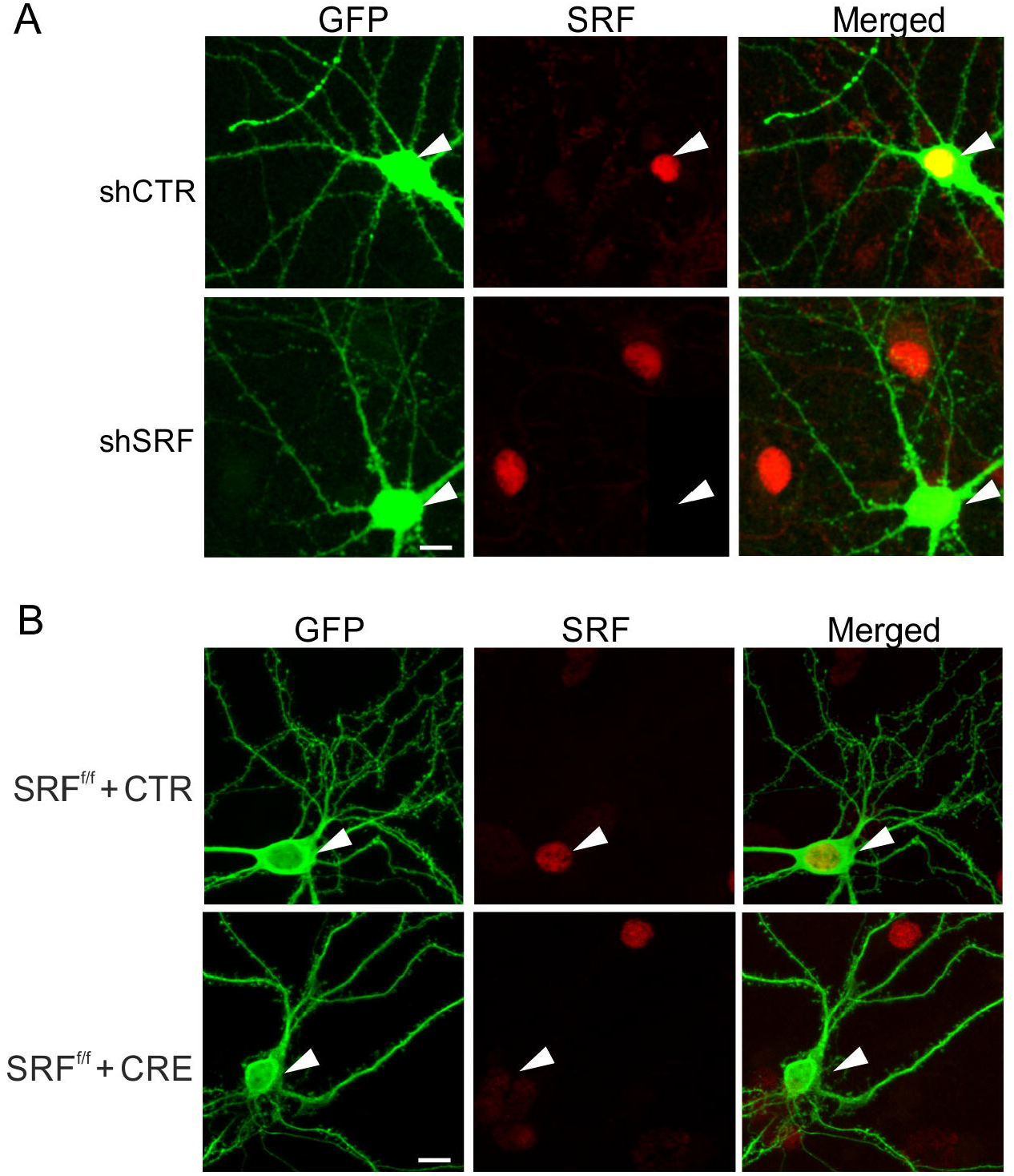
SRF knockdown in primary hippocampal neurons reduced SRF expression *in vitro*. **(A, B)** Representative images of **(A)** GFP-positive rat hippocampal neurons that were transfected with shCTR or shSRF plasmid and (**B**) SRF^f/f^ mouse hippocampal neurons that were co-transfected with CaMKIIα-GFP (control) together CaMKII-Cre-recombinase or CaMKIIα-GFP alone and immunostained with anti-SRF antibody (red). The white arrowheads indicate transfected cells. Scale bar = 10 μm.

**Fig. S2.**
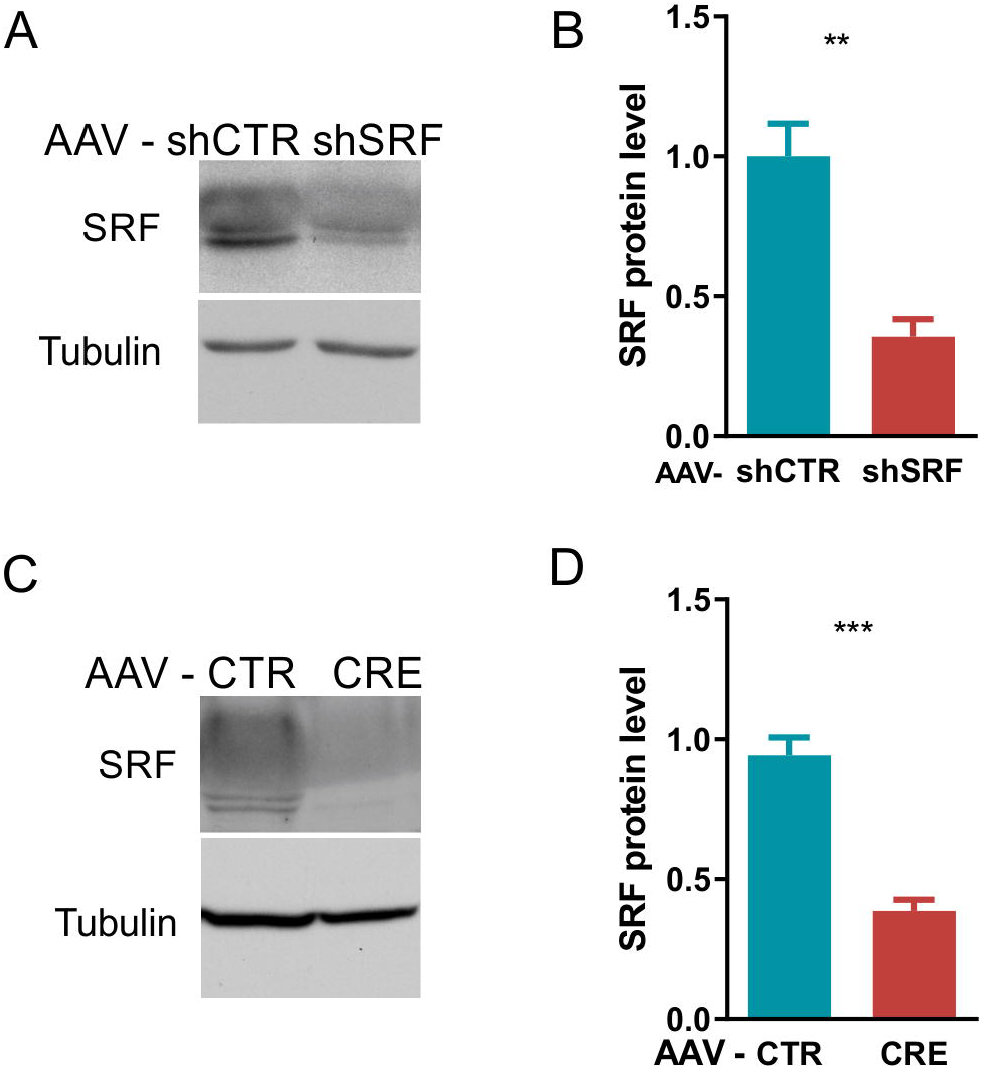
AAV-mediated transduction of primary hippocampal neurons downregulates SRF expression. **(A, B)** Western blots of protein extracts from **(A)** AAV-shCTR or AAV-shSRF-transduced rat hippocampal neurons and **(B)** AAV-CAMKIIα-mCherry or AAV-CAMKIIα-Cre-transduced *Srf*^f/f^ mouse hippocampal neurons. **(D)** Quantification of SRF total protein levels in AAV-shCTR- and AAV-shSRF transduced-rat hippocampal neurons. (**E**) Quantification of SRF total protein levels in AAV-CAMKII-mCherry- and AAV-CAMKIIα-Cre - transduced *Srf*^f/f^ mouse hippocampal neurons. Tubulin served as the loading control. *\*\*p* < 0.01, *\*\*\*p* < 0.001 (Mann-Whitney test). All of the data are presented as a fold change relative to control. The data are from at least three independent neuronal cultures and expressed as mean ± SEM. The data from rat hippocampal cultures in **(B)** passed the normality test. \*\**p* < 0.01 (Student’s *t*-test).

**Fig. S3.**
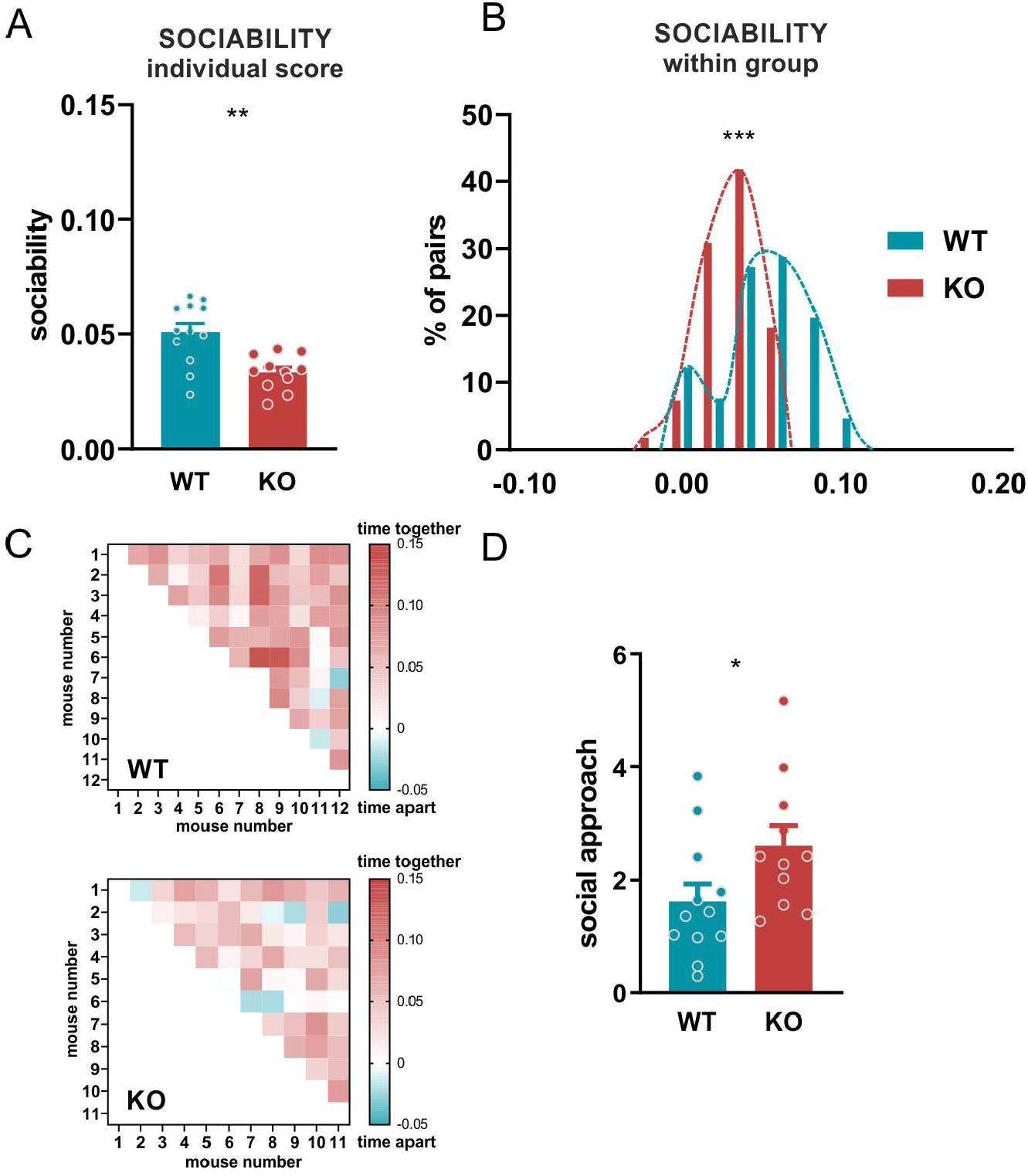
SRF KO female mice exhibit decrease in sociability, similar to males. **(A)** Sociability, measured as the time voluntarily spent together with others, in WT and SRF KO mice. **(B)** Histogram of the distribution of in-cohort sociability in WT and SRF KO mice. \*\*\**p* < 0.0001 (Kolmogorov-Smirnov test). **(C)** Raw data from **(A)** and **(B)**, represented as a matrix in which squares illustrate the time voluntarily spent together by each pair of mice within the cohort. The intensity of colors reflects the strength of the relationship in accordance with the presented scale. **(D)** Social approach in WT and SRF KO mice. *n* = 12 female WT mice; *n* = 11 female SRF KO mice. \**p* < 0.05, \*\**p* < 0.01, \*\*\**p* < 0.001 in **(A)** and **(D)**(Mann-Whitney test). The data are expressed as mean ± SEM. Sociability and social approach data in **(A)** and **(D)** passed the normality test. \**p* < 0.05, \*\**p* < 0.01 (Student’s *t*-test).

## Notes

### Competing Interest Statement

The authors have declared no competing interest.

